# A likelihood ratio test for changes in homeolog expression bias

**DOI:** 10.1101/119438

**Authors:** Ronald D. Smith, Taliesin J. Kinser, Gregory D. Conradi Smith, Joshua R. Puzey

## Abstract

**Background:** Gene duplications are a major source of raw material for evolution and a likely contributor to the diversity of life on earth. Duplicate genes (i.e., homeologs, in the case of a whole genome duplication) may retain their ancestral function, sub- or neofunctionalize, or be lost entirely. A primary way that duplicate genes may evolve new functions is by altering their expression patterns. Comparing the expression patterns of duplicate genes may give clues as to whether any of these evolutionary processes have occurred.

**Results:** We develop a likelihood ratio test for the analysis of the expression ratios of duplicate genes across two conditions (e.g., tissues). We demonstrate an application of this test by comparing homeolog expression patterns of 1,448 homeologous gene pairs using RNA-seq data generated from the leaves and petals of an allotetraploid monkeyflower *(Mimulus luteus*). We assess the sensitivity of this test to different levels of homeolog expression bias and compare the method to several alternatives.

**Conclusions:** The likelihood ratio test derived here is a direct, transparent, and easily implemented method for detecting changes in homeolog expression bias that outperforms three alternative approaches. While our method was derived with homeolog analysis in mind, this method can be used to analyze changes in the ratio of expression levels between any two genes in any two conditions.

## Background

Gene duplications are a major source of raw material for evolution and a likely contributor to the diversity of life on earth [1–9]. Gene duplications are a special type of mutation resulting in the multiplication of intact functional components. These duplicate genes may either retain the ancestral function or individual portions of the gene’s ancestral function may be partitioned (i.e., subfunctionalize) or evolve new functions entirely (i.e., neofunctionalize) [10–12]. Duplicate genes may evolve new functions either by changes in the primary coding sequence or altering where and when they are expressed. Previous work has indicated that changes to gene expression and their regulatory networks may be more important, rapid, or flexible than divergence of protein identities in the evolution of sub- and neofunctionlization [13–19].

There are multiple scenarios in which genes can be duplicated, ranging from small regional gene duplications to massive whole genome duplications (WGDs).

The term polyploid refers to cells or organisms that have undergone a WGD event and contain more than two paired sets of chromosomes. Each complete set of chromosomes is referred to as a subgenome. Homologous genes located on separate subgenomes are referred to as homeologs.

WGDs are especially common in plants; indeed, all extant angiosperms (i.e., flowering plants) have at least two rounds of WGD in common [20] and up to 15% of speciation events in angiosperms may have been the product of WGDs [21]. Importantly, all major crops (rice, corn, potato, wheat, etc.) are polyploid [22]. WGD events and the resulting polyploidy are not restricted to plants, but have occurred in both vertebrate and invertebrate lineages as well. For example, the African clawed frog, *Xenopus,* commonly used as an experimental model system and extensively studied in developmental biology, includes species ranging from diploid to dodecaploid [23]. Other examples of polyploids with ancient WGD events include the the zebrafish *Danio rerio* [24], several salmonids [2], and some species of fungi [25]. Interestingly, there exists at least one polyploid mammal [26], a tetraploid rat from Argentina that mediates gene dosage by regulation of ribosomal RNA.

The biological consequences of gene duplications and subfunctionalization are significant and include examples such as the evolution of eyes [27], the evolution of hemoglobins [28], development of heat resistance in plants [29], and insecticide resistance [30]. Given the importance of duplicate genes in evolution, it is natural to ask how we might quantify differences in the activity or function of homeologous genes. One way to begin exploring this question is by analyzing gene expression levels.

Genome-wide gene expression levels are commonly quantified using high throughput RNA sequencing (RNA-seq) [31]. In RNA-seq experiments, mRNA is extracted, purified, and reverse transcribed into cDNA. This cDNA is fragmented into smaller pieces and sequenced using next-generation technology. The resulting millions of sequence reads are then mapped to either a reference genome or reference transcriptome, and the number of sequences mapping to a particular gene is used as an indication of the expression level of that gene.

In *differential expression analysis,* high-throughput RNA-seq data is used to determine if gene expression levels vary under different experimental conditions, or in distinct tissues, etc. Several different approaches to this statistical analysis exist [32–34], some of which use methods based on maximum likelihood estimation and likelihood ratio tests.

Homeologous gene pairs frequently have distinguishing sequence differences. Therefore, sequencing reads derived from individual homeologs can be distinguished and expression levels can be determined for each homeolog. The term *homeolog expression bias* (HEB) refers to cases where homeologs are expressed at unequal levels in a single experimental condition [35]. The primary objective of this paper, development of a likelihood ratio test for statistical analysis of *changes* in homeolog expression bias (denoted ΔHEB) is a non-trivial extension of the statistical analysis of differential expression.

The following sections begin with the derivation of a likelihood ratio test for HEB. This is our starting point for the development of a likelihood ratio test for ΔHEB, i.e. changes in relative expression levels between homeologous genes in two conditions. We demonstrate an application of this method using RNA-seq data to compare homeologous gene expression in petals and leaves of the allotetraploid *Mimulus luteus*. Finally, using simulated data, we show that the likelihood ratio test for ΔHEB derived here is the best choice among competing methods.

## Methods

### Quantifying homeolog expression bias (HEB)

We will write *A* and *B* to denote a homeologous gene pair from which RNA-seq data is generated in *n* biological replicates. Typically, the mean expression levels of the homeologs (denoted *a̅* and *b̅*) are normalized by gene length and sequencing depth, as when reported in units of RPKM (reads per kilobase of coding sequence per million mapped reads). We define the homeolog expression bias (HEB) of the *n* replicates as

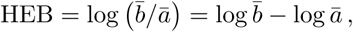

a dimensionless quantity with HEB = 0 indicating no bias. If one uses the base 2 logarithm, HEB = −3 indicates 8-fold bias towards homeolog *A*.

### Likelihood ratio test for HEB

After accounting for the possibility of different gene lengths, the statistical test for HEB is essentially a likelihood ratio test for differential expression of a pair of homeologous genes. The goal is to determine whether there is sufficient evidence to reject the null hypothesis (*H*_0_) that there is no bias (i.e., equal expression levels for homeologous genes) in favor of the alternative hypothesis (*H*_1_) that bias is present, i.e., different expression levels for homeologous genes. In mathematical terms, the null hypothesis *H*_1_ corresponds to the parameters (denoted by *θ*) of a probability model for generating the data being in a specified subset Θ_0_ of the parameter space Θ, that is,

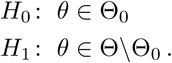

Let θ = (λ^*a*^, λ^*b*^) denote the true but unknown expression levels (properly scaled, e.g., in units of RPKM). Assuming positive, i.e. non-zero, expression, the parameter space is Θ = {θ: λ^*a*^, λ^*b*^ ∈ ℝ_+_}. The null (*H*_0_) and alternative (*H*_1_) hypotheses for the likelihood ratio test for homeolog expression bias are formalized as follows,

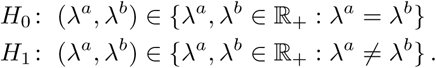

Equivalently, let *ω* = λ^*b*^/λ^*a*^ denote the ratio of expression levels and drop the superscript indicating the reference homeolog (λ = λ^*a*^). In that case, λ^*b*^ = *ω*λ and the hypotheses are written as follows,

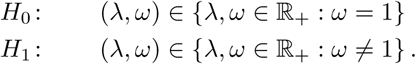

Once we specify a probability model for the data χ, likelihood functions for each hypothesis, 𝓛_0_(*θ*|χ) and 𝓛_1_(*θ*|χ), can be derived (see next section). For composite hypotheses, the appropriate likelihood ratio test statistic is

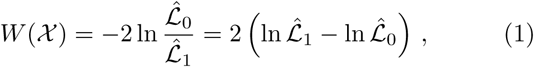

where 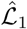 are the maximized likelihoods,

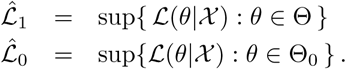

A critical value of the test statistic (*W*_*_) is obtained from the Chi-squared distribution with significance level *α* = 0.05. The number of degrees of freedom *δ* is the difference in the number of free parameters in Θ and Θ_0_ (here *δ* = 1) [36]. The null hypothesis *H*_0_ is rejected in favor of the alternative *H*_1_ when *W*(χ) > *W*_*_.

### Probability model for RNA-seq read counts

Denote the lengths of homeologous genes *a* and *b* as *ℓ^*a*^* and *ℓ^b^* (e.g., in kilobases) and let *d_i_* be the sequencing depth (e.g., in millions of mapped reads) of replicate *i*. The expected number of RNA-seq reads for gene *a* and replicate *i* is

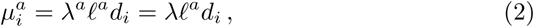

where in the second equality we have dropped the superscript for the reference homeolog (λ = λ^*a*^). Similarly, the expected number of RNA-seq reads for gene *b* and replicate *i* is

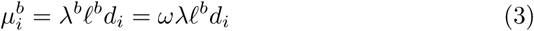

where *ω* = λ^*b*^/λ^*a*^ = λ^*b*^/λ.

The probability model assumes that the count data for each gene is drawn from a negative binomial distribution,

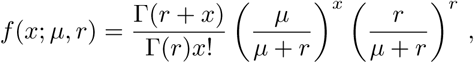

where *μ* is the appropriate mean (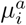 in Eqs. 2 and 3). That is, if 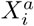 are random variables representing the count data for replicate *i* of homeologous genes A and B,

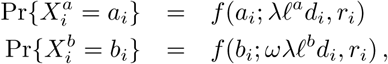

where we have used 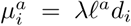 In these expressions, the aggregation parameter *r_i_* is obtained from the observed mean-variance relation for all homeolog pairs of the ith experimental replicate (see Appendix 1).

Assuming independence of experimental replicates, the likelihood functions 𝓛_1_ and 𝓛_0_ are products of the likelihood functions for each observation, that is,

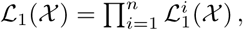

and similarly for 𝓛_0_(χ), where χ_*i*_ = {*a_i_*, *b_i_*} indicates the observed read counts for replicate *i* and 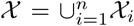 The likelihood function for the alterna-tive hypothesis and the ith replicate is

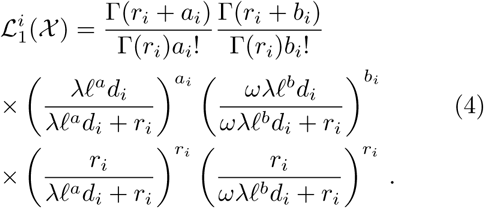

The likelihood function for the null hypothesis and the *i*th replicate, 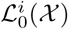 is given by Eq. 4 with *ω* = 1.

### Maximum likelihood estimation

Maximum likelihood estimation is performed using the the log-likelihood function corresponding to Eq. 4, namely,

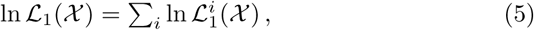

where

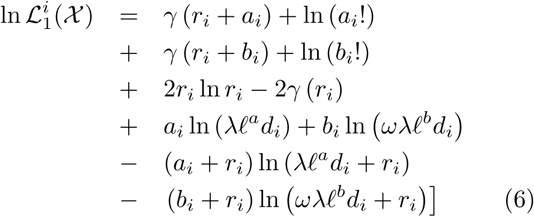

and γ(·) = lnΓ(·). The log-likelihood function for the null hypothesis (ln 𝓛_0_) is given by Eq. 13 with *ω* = 1.

The log-likelihood function ln L1(X) is maximized by numerically solving for 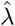 leading to zero partial derivatives,

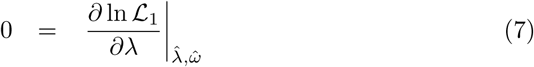

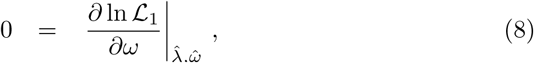

as described in Appendix 2. The log-likelihood function ln 𝓛_0_(χ) is maximized by solving for λ̂ leading to

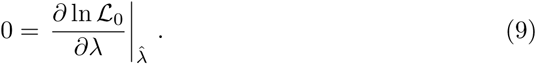

The optimal parameter values 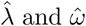 are used to evaluate ln 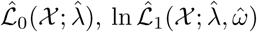 and the test statistic *W* (see Eq. 1).

### Quantifying changes in homeolog expression bias (△HEB)

Let *A* and *B* represent homeologous genes and RNA-seq data is generated under conditions 1 and 2 in n biological replicates, leading to mean expression levels *a̅*_1_, *a̅*_2_, *b̅*_1_, *b̅*_2_. The change in homeolog expression bias (ΔHEB) is defined as

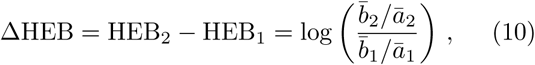

where the last equality uses HEB_1_ = log *b̅*_1_/*a̅*_1_ and HEB_2_ = log *b̅*_2_/*a̅*_2_.

### Likelihood ratio test for ΔHEB

The likelihood ratio test for ΔHEB is designed to determine whether there is sufficient evidence to reject the null hypothesis (*H*_0_) that homeolog expression bias is the same under two experimental conditions (ΔHEB = 0) in favor of the alternative hypothesis (*H*_1_) that there is a difference in bias)( △HEB ≠ 0). Following notation similar to the previous section, our hypotheses are

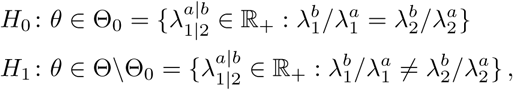

where 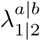 is an abbreviation for 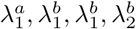 Equivalently,

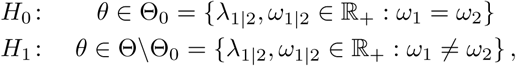

where 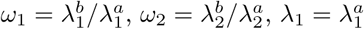 The difference in degrees of freedom of the alternative and null hypotheses is *δ* = 4 − 3=1.

The likelihood functions for the ΔHEB test are similar to those for HEB, though the two different experimental conditions lead to twice as many terms (cf. Eq. 4). The likelihood function for *H*_1_ is

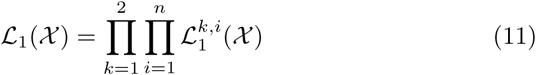

where 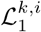 the likelihood function for the *i*th replicate of the *k*th condition, has the form of Eq. 4 with parameters indexed by condition 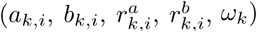 The log-likelihood function for *H*_1_ is thus

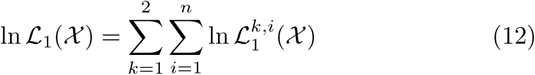

where

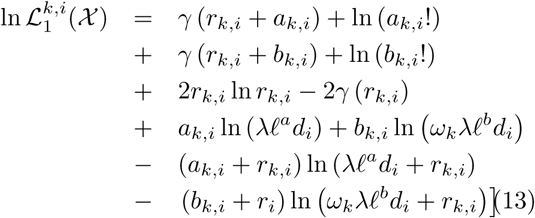

and γ(·) = lnГ(·). The log-likelihood function for the null hypothesis (ln 𝓛_0_) is given by the above expressions with ω_1_ = ω_2_ = ω. The aggregation parameters (*r_k, i_*) are determined from the data with experimental conditions *k* = 1 and 2 considered separately (cf. Eqs. 17–19).

The log-likelihood function ln 𝓛_1_(χ) used in the analysis of ΔHEB is maximized by numerically solving uncoupled systems of the form of Eqs. 7 and 8 for 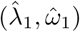 The log-likelihood function ln 𝓛_0_(χ) is maximized by solving for 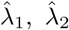 that lead to zero partial derivatives,

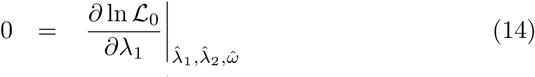

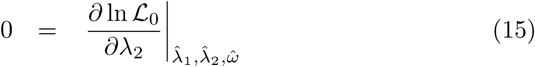

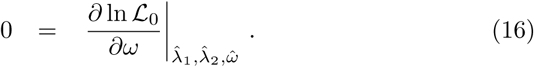

The optimal parameter values are used to evaluate the likelihoods, 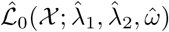 and the test statistic W (see Eq. 1).

The numerical solution of these equations was facilitated by transforming these equations in a manner that ensured both parameters are positive and symmetric with respect to the mean expression levels of homeolog *A* and *B* (see Appendix 2).

## Results

### The likelihood ratio test for HEB applied to allotetraploid *Mimulus luteus*

To demonstrate the application of the likelihood ratio test for HEB, five biological replicates of RNA-seq data were generated from petals of the tetraploid *Mimulus luteus* (monkeyflower), and another five replicates were generated from the leaves (see Appendix 3 for details). We have chosen *M. luteus* because it is a tetraploid with two distinct subgenomes [37], denoted A and B. In this section, we use the likelihood ratio test for HEB to find homeologous gene pairs where one homeolog is expressed at significantly different levels than the other, one tissue at a time. In the section on ΔHEB we develop a likelihood ratio test to determine whether there is a significant difference in the bias between the two tissues.

#### Homeolog expression bias in Mimulus luteus petals

Figure 1 (top panel) shows the result of applying the likelihood ratio test for HEB to the petal data. There are 1,853 homeologous gene pairs in *M. luteus* that can be identified as coming from separate subgenomes. Of these 1,853 homoeologous pairs, 1,560 were testable (measurable expression from each individual homeolog). Of the testable pairs, a total of 676 gene pairs show significant bias (using a significance level of *α* = 0. 05, and applying the Benjamini-Hochberg correction [38, 39] to account for multiple testing error). In the 334 pairs biased towards the *A* homeolog the mean HEB is −2.49 (5.6-fold change). In the 342 pairs biased towards the *B* homeolog, the mean HEB is 2.39 (5.2-fold change).

**Figure 1:**
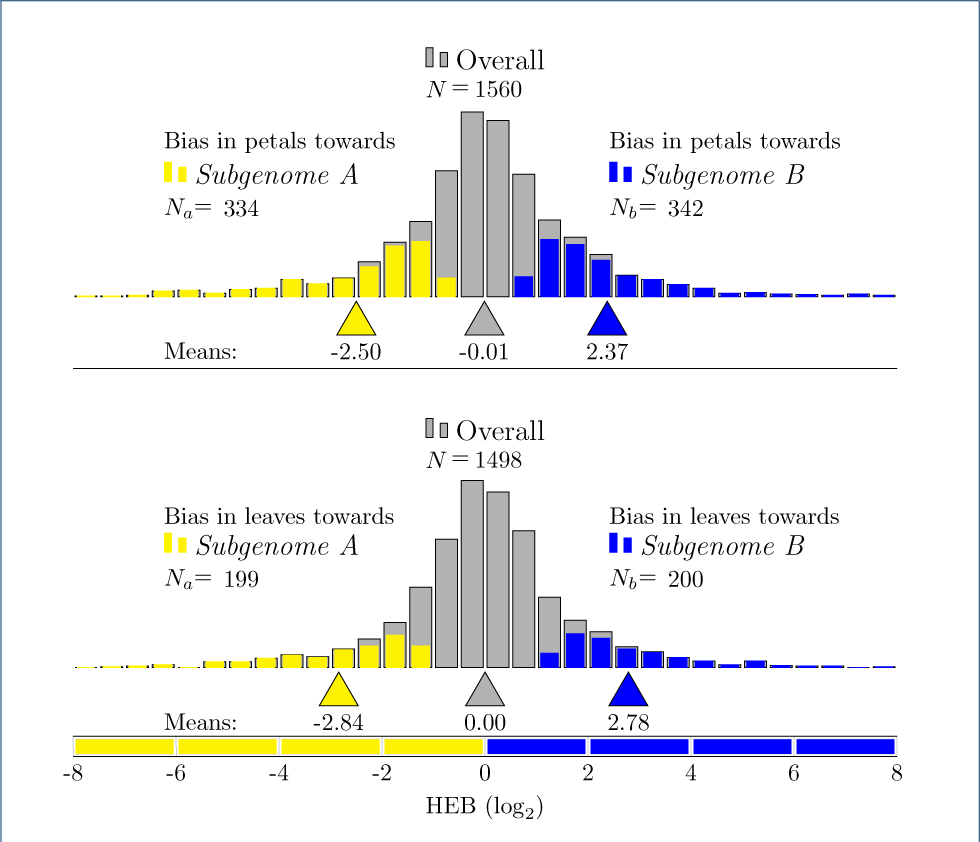
Likelihood ratio test for HEB in petals (top) and leaves (bottom) of M. *luteus*. (**Top**) Of 1,560 testable homeologous gene pairs in the petals (gray), a total of 676 show significant bias. Of these, 334 pairs are biased towards the *A* homeolog (yellow), with a mean HEB of −2.49 (5.6×). 342 pairs are biased towards the *B* homeolog (blue), with a mean HEB of 2.39 (about 5.2×). (**Bottom**) Of 1,560 testable homeologous gene pairs (gray), a total of 676 show significant bias. Of these, 334 pairs are biased towards the *A* homeolog (yellow), with a mean HEB of −2.49 (5.6×). 342 pairs are biased towards the *B* homeolog (blue), with a mean HEB of 2.39 (about 5.2×). The Benajamini-Hochberg correction for multiple testing was applied at significance level *α* = 0.05 (and also in Figure 3).

These results may be indicative of a number of evolutionary processes. For example, one of the homeologs may have become sub- or neofunctionalized in this tissue, or one of the homeologs may simply be losing its function.

#### Homeolog expression bias in Mimulus luteus petals

Next, the likelihood ratio test for HEB was applied tothe leaf data (results shown in Fig 1, bottom panel). Ofthe 1,853 homoeologous pairs, 1,498 were testable. Ofthis subset, a total of 399 gene pairs show signicant bias. In the 199 pairs biased towards the *A* homeolog the mean HEB is −2.83 (7.1-fold change). In the 200 pairs biased towards the *B* homeolog, the mean HEB is 2.80 (7.0-fold change).

### The likelihood ratio test for ΔHEB applied to allotetraploid Mimulus luteus

The likelihood ratio test for ΔHEB requires each homeolog to have at least one read in each condition. Returning to the leaf and petal data from the previous sections on HEB, this gives 1,448 testable pairs. Figure 2 shows the results of the likelihood ratio test for ΔHEB. We find a total of 76 gene pairs show significant ΔHEB. Of these, 35 are more biased towards the *A* homeolog in the leaf than they are in the petal. The remaining 41 gene pairs are more biased towards the *B* homeolog in the leaf than they are in the petal.

**Figure 2:**
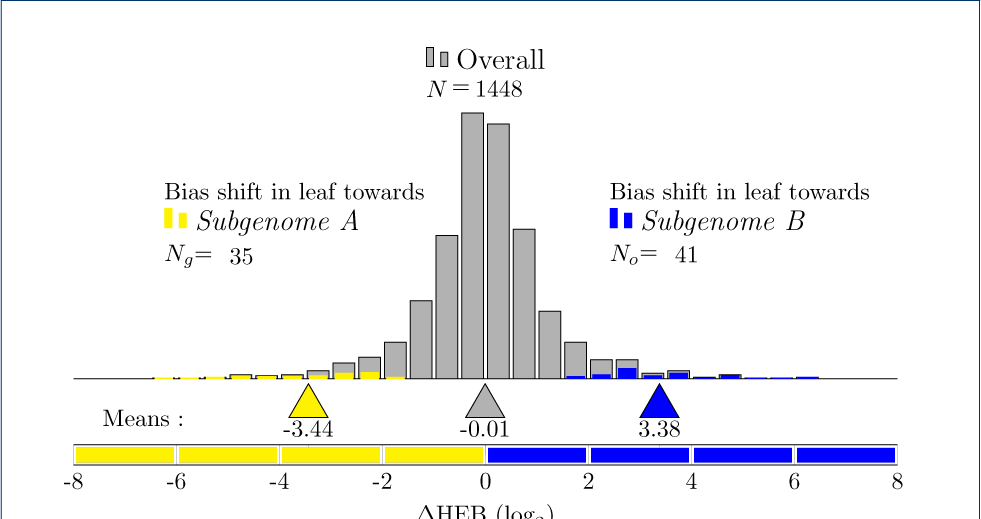
Likelihood ratio test for ΔHEB in the leaves vs. petals of *M. luteus*. Of 1,448 testable homeologous gene pairs (gray), 76 show significant △HEB. Of these, 35 are more biased towards the *A* homeolog in the leaves than in the petals (yellow). 41 gene pairs are more biased towards the *B* homeolog in the leaf than in the petal (blue).

Figure 3 shows a scatter plot of homeolog expression bias (HEB) in leaf and petal. Colored marks indicate gene pairs with statistically significant changes in homeolog expression bias ΔHEB) (these points correspond to the colored bars in Figure 2). Data points in the top-left and bottom-right quadrants of Figure 3 represent homeologous pairs where one homeolog is more highly expressed in one tissue and its partner is more highly expressed in the other tissue. On the other hand, the top-right and bottom-left quadrants correspond to homeologous pairs where the difference in bias favors the same homeolog but has become more extreme. Finally, all of the marks that are colored blue or yellow show significant change in bias and are candidates for tissue specific sub- or neofunctionalization.

**Figure 3:**
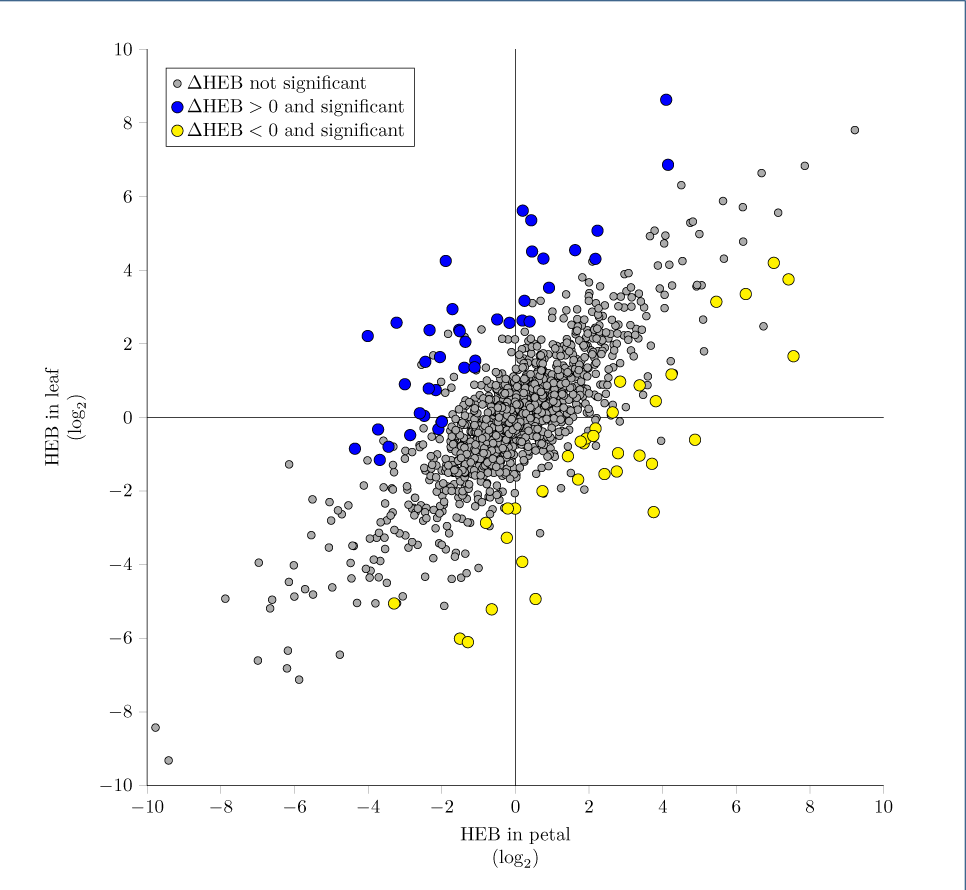
Statistical significance of ΔHEB. Compared to homeolog expression bias (HEB) in leaf and petal. Yellow and blue indicates homeolog gene pairs with significant AHEB. The likelihood ratio test for AHEB is distinct from HEB tests in leaf and petal (see text).

Although the change in homeolog expression bias is defined by Eq. 10 as the log-fold change in homeolog expression bias, the intercalation of significant (yellow and blue) and not significant (gray) ΔHEB in Figure 3 makes it clear that statistical evidence for ΔHEB is not reducible to the difference between HEB_leaf_ and HEB_petal_ (the vertical or horizontal distance to the line of slope 1 where HEB_leaf_ = HEB_petal_).

Whether or not ΔHEB can be called significant also depends on differences in sequencing depths, mean expression levels (e.g., lowly expressed genes are more likely to be influenced by shot noise), and ratios of gene lengths. All of these factors are considered simultaneously in the likelihood ratio test presented here. Calling ΔHEB based on sequential HEB results would almost certainly result in a different set of genes being called significant.

### Validation of the likelihood ratio tests using simulated data

A natural question to ask about HEB and ΔHEB is, “How large does the change in expression levels between homeologs across conditions need to be before we can detect ΔHEB most of the time?”. Unsurprisingly, this depends largely on the number of biological replicates.

To explore this question we generated simulated data with one expression level fixed at a constant value, μ^*a*^ = 100, and varied the other expression level, μ^*b*^ = 2^*x*^ *μ^a^* with *x* ∈ [0, 2] in steps of 0.1. For each value of x, we generated 10,000 sets of data from a negative binomial distribution for *N* = 3, 6,12 and 24 replicates. We fixed the parameter *r* = 10 for simplicity; this is in the typical range of values we have observed in RNA-seq data.

Fig 4 shows the results of the likelihood ratio test for HEB on this simulated data set. We find that a 4fold change is almost always detectable, regardless of the number of replicates. However, detecting a 2-fold change at least 95% of the time requires at least 12 replicates.

**Figure 4:**
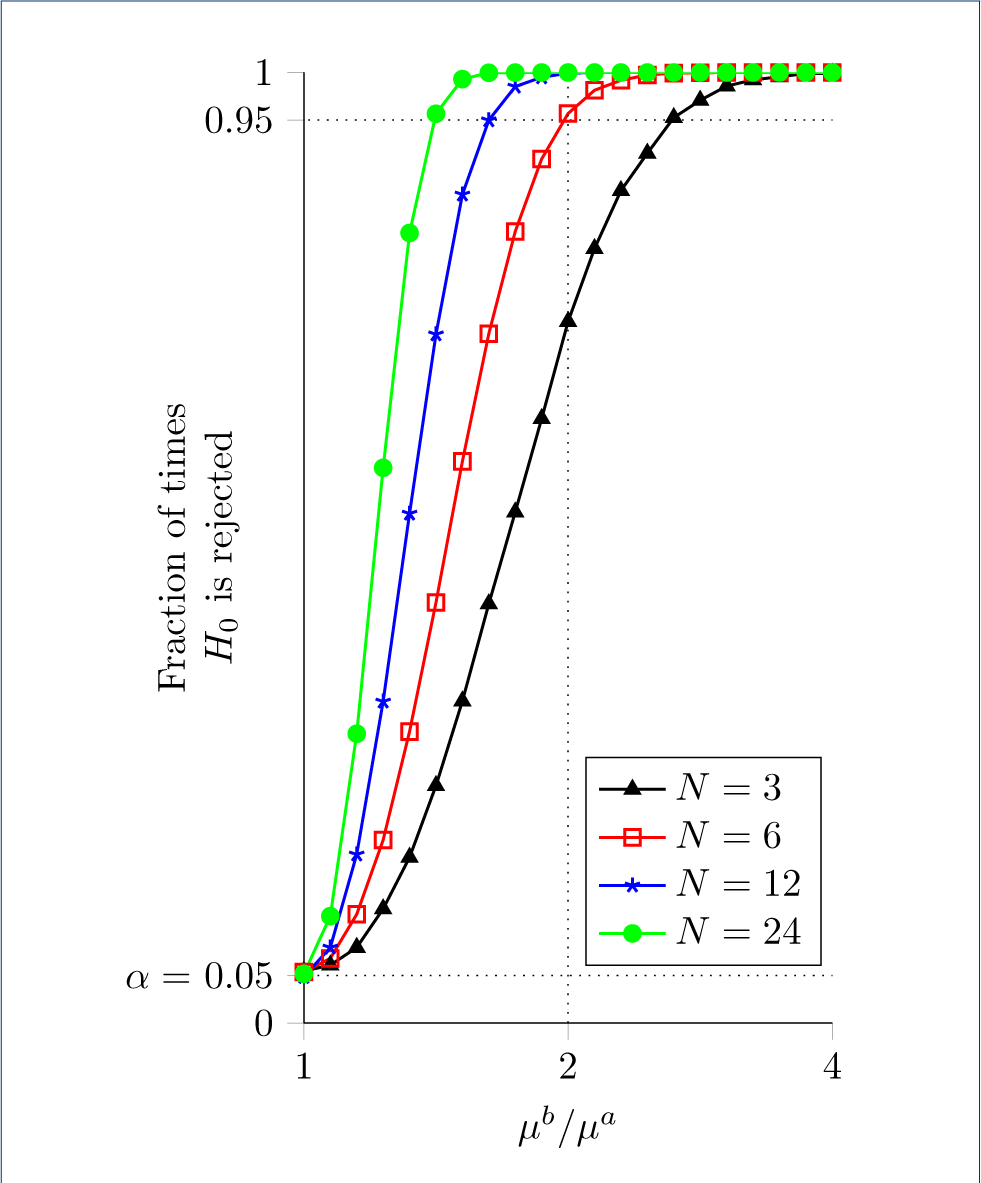
Ability of the likelihood ratio test for HEB to detect different levels of bias. Simulation results show the fraction of times *H*_0_ was rejected for 10,000 trials with the given values of *x* and *n* (parameters: *α* = 0.05, *μ*^*a*^ = 100, *r* = 10). With *n* ≥ 3 replicates, a 4-fold change is detectable over 95% of the time. Detecting a 2-fold change greater than 95% of the time requires at least 12 replicates.

To assess the sensitivity of ΔHEB to different levels of bias shift, we created a similar data set. This time, we set 3 of the expression levels equal 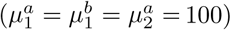 100), and varied the fourth; 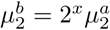 in steps of 0.1. The aggregation parameter was again fixed at *r* = 10. For each value of *x*, 10,000 sets of data were generated from a negative binomial distribution for *N* = 3, 6,12 and 24 replicates.

Fig 5 shows the results of the likelihood ratio test for ΔHEB on this simulated data set. The results are similar to those for HEB, with the test for ΔHEB being slightly less sensitive than the test for HEB. For ΔHEB, a 4-fold change in bias is detected more than 95% of the time when N ≥ 6. As with the test for HEB, the ability to detect smaller changes increases significantly with the number of replicates.

**Figure 5:**
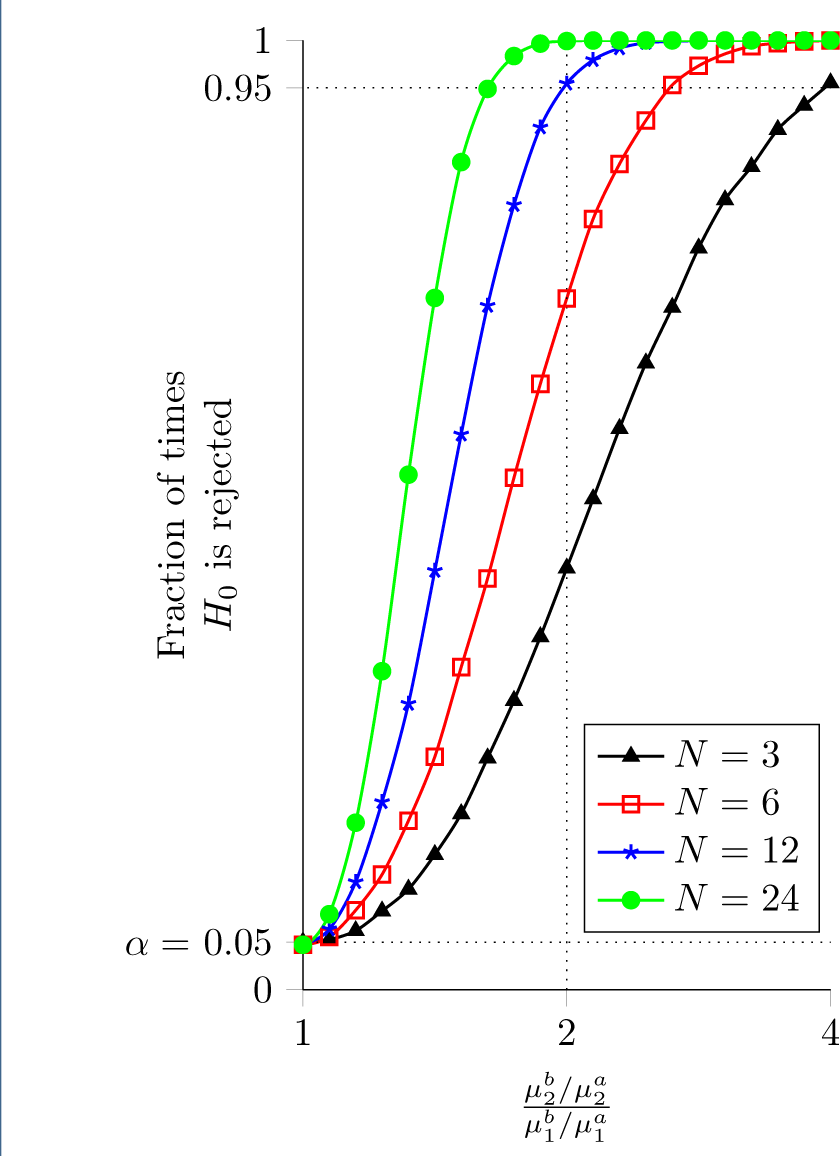
Ability of the likelihood ratio test for ΔHEB to detect different levels of change in bias. Simulation results show the fraction of times *H*_0_ was rejected for 10,000 trials with given values of *x* and *n* (parameters: *α* = 0.05, 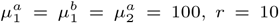 With *n* ≥ 6 replicates, a 4-fold change is detectable over 95% of the time. However, detecting a 2-fold change more than 95% of the time requires at least 24 replicates.

## Discussion

### Alternative Methods

While our method is transparent, derived specifically for the analysis of ΔHEB, and requires a minimal number of assumptions, we also wished to investigate whether other methods could achieve similar results. Since we found nothing directly comparable to our method in the literature, we developed three additional ad hoc methods. To compare these methods we generated simulated data sets and analyzed ROC curves. Each data set contained 20,000 gene pairs, half of which had ΔHEB fixed at a constant value (2,8, and 16). Three replicates were generated from negative binomial distributions, and this was repeated 100 times for each value of ΔHEB (300 simulations total).

First, we took a naive approach and performed t-tests and z-tests on the ratio of log2 -fold changes between conditions 1 and 2. Next, we ran DESeq2 and extracted the estimated shrunken log2 -fold changes and their standard errors, and performed a z-test (we call this method ‘DEZ’). Unsurprisingly, the naive methods underperformed the LRT, with area under the ROC curve (ROC area) typically less than the LRT by « 0.05 to 0.36.

The LRT outperformed DEZ for ΔHEB = 8 and 16 (Figure 6, top region). For ΔHEB = 2, both methods performed poorly with mean ROC area = 0.5616 for the LRT, while DEZ came out slightly ahead with mean ROC area = 0.5620 (not shown).

**Figure 6:**
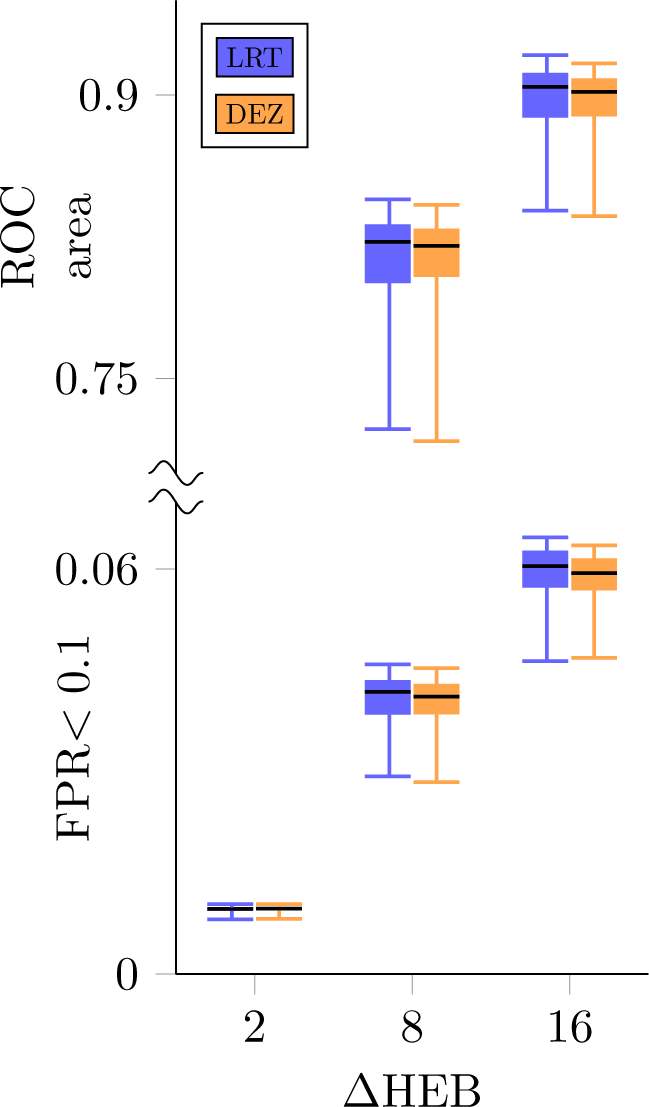
Comparison of the LRT and DEZ. The top part of the plot shows the distribution of area under ROC curves for 100 trials of simulated data. Each trial contained 20,000 genes, half of which had △HEB fixed at a constant value. Results for △HEB = 2 are not shown as they were too low (mean area for LRT= 0.5616, for DEZ=0.5620). The bottom part of the plot shows ROC area constrained to false positive rates less than 0.1. In both regions, boxes indicate interquartile ranges, whiskers indicate 5th and 95th percentiles, and black lines indicate medians.

It is possible for a test to have a larger ROC area but not necessarily be the best choice, for example, if a curve accumulates a small amount of area for low FPR, and a large amount of area for high FPR. To address this we evaluated partial ROC area for false positive rates between 0 and 0.1 (we assume that, in practice, most researchers would never accept FPR> 0.1). By this metric, the LRT outperforms DEZ for ΔHEB = 8 and 16, while for ΔHEB = 2 both methods performed poorly, with DEZ marginally better (Figure 6, bottom region).

## Conclusion

Gene duplication and polyploidy are extremely important factors in generating the diversity of life on earth. As Ohno stated in his seminal work on gene duplication [1], “Natural selection merely modified while redundancy created” the raw materials necessary for the diversification of life on earth.

In this paper we have developed a robust statistical framework specifically designed for the comparison of duplicate gene expression patterns. Importantly, this technique is consistent and reproducible. Through analysis of simulated data we have shown that these methods perform well, especially given the typically small sample sizes in most RNA-seq experiments. We have shown that the ability to detect small differences in expression levels increases as a function of sample size, a fact which can be used to aid experimental design. Other authors have noted this with traditional differential expression analysis and made similar recommendations [40–42]. Moreover, we demonstrate the usefulness of the likelihood ratio test for ΔHEB using homeolog expression (RNA-seq) data derived from a polyploid plant. While we have developed this test for the purpose of analyzing changes in expression patterns of homeologous genes, we emphasize that the methods are suitable for the expression analysis of any two genes (they need not be homeologs) across any two conditions.

## APPENDIX 1: Estimation of aggregation parameters

Due to the typically small number of replicates in RNA-seq experiments, accurate estimation of the aggregation parameter is not realistic on a gene-by-gene basis [34, 43]. Instead, we use the mean-variance relation of a negative binomial distribution, namely,

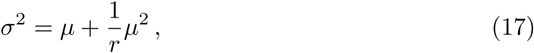

to compute an aggregation parameter *r* for each experimental replicate, after rescaling to account for each replicates sequencing depth.

In brief, let 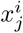 denote the count data for the *j*th pair of homeologous genes obtained for experimental replicate *i* ∈ {1, 2, . . ., *n*}. For each of the *n* replicates, we produce an auxiliary data set 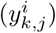 by rescaling the count data for all replicates as though each were obtained in an experiment with the sequencing depth of replicate *k*,

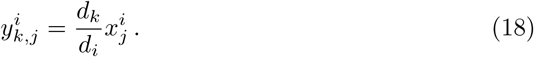

For each gene (*j*), we compute a scaled mean (μ_*k,j*_) and variance 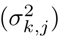 over replicates (*i*). To obtain the aggregation parameter *r_k_*, we perform a nonlinear least squares fit of the observed mean-variance relation across all genes. That is, *r_k_* minimizes the sum of squares error,

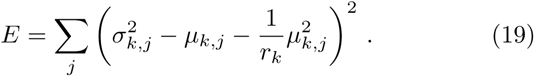

## APPENDIX 2: Numerical scheme for maximum likelihood estimation

For the analysis of both HEB and ΔHEB, parameter values maximizing the likelihood functions 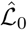 were obtained using the built-in MATLAB command `fsolve` applied to Eqs. 7–9 and 14–16. In both cases, the numerical procedure was facilitated by changing variables from (λ, ω) to (υ, *y*) through

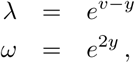

that is, υ = ln λ + *y* and *y* = (ln ω)/2. This ensures positivity of Λ and ω and leads to a system of equations that is symmetric in λ^*a*^ ↔ λ^6^. The new variable v is the logarithm of the geometric mean of the expression levels λ^*a*^ = λ and λ^*b*^ = ω λ,

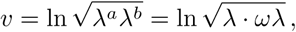

that is, λ^*a*^ = λ = *e^υ–y^* and λ^*b*^ = ω λ = *e*^υ+*y*^. The transformed partial derivatives used to maximize the log-likelihood ln 𝓛_1_ (Eqs. 7–8) are

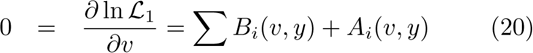

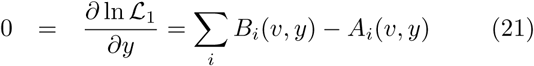

where

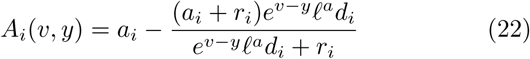

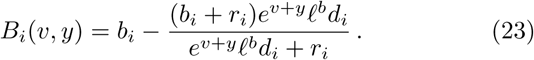

The transformed partial derivative used to maximize ln 𝓛_0_ are found by substituting *y* = 0 in Eq. 20,

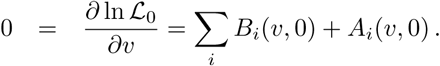

For the analysis of ΔHEB, the partial derivatives used to maximize ln 𝓛_1_ are two uncoupled systems of the form of Eq. 20–23, one for each experimental condition (*k* = 1 and 2),

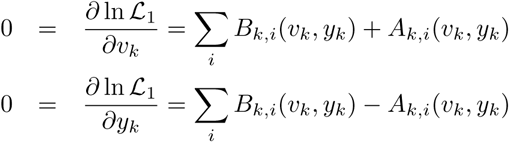

where

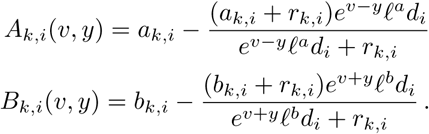

For the null hypothesis *y*_2_ = *y*_1_ = *y* we numerically solve a system of three equations, including

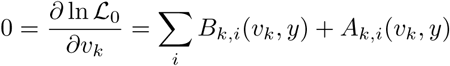

for *k* = 1 and 2. These are coupled via

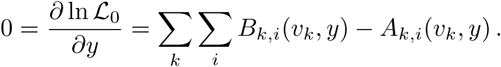

## APPENDIX 3: Experimental methods

Plant tissues were collected from second generation inbred *Mimulus luteus.* All plants were grown in a greenhouse under a 16 hour light regiment at 21° C and 30% humidity. Petal tissue was collected from the corolla of a flower bud near blooming, and leaf tissue came from young leaves adjacent to the stem apical meristem. Five replicates of each tissue type were collected, at the same time of day, from different individuals. Approximately 100-200 mg of plant tissue was immediately placed into liquid nitrogen. RNA was extracted by grinding frozen tissue with pestles in PureLink© Plant RNA Reagent from Ambion^TM^. Column isolation of RNA was subsequently performed using Direct-zol^TM^ RNA MiniPrep Plus Kit from Zymo Research. Libraries were constructed using KAPA Stranded mRNA-Seq Kit. During library construction, sequence specific Illumina TruSeq©R adapters were added to distinguish each library. Using an Agilent 2100 Bioanalyzer, average fragment lengths were determined to be between 230 and 300 bp. Libraries were then pooled and sequenced by the Duke Center for Genomic and Computational Biology on an Illumina HiSeq 2500 instrument. The resulting reads (50 base pair, single end) were mapped to the *M. luteus* genome using bowtie2 [44] with the `--very-sensitive-local` option. Reads to exonic regions were counted using `htseq-count` [45] with the default settings (minimum alignment quality of 10 on the phred scale).

## Availability of data and materials

Sample MATLAB code and the data used in ΔHEB analysis can be found on the Mathworks file exchange, submission number 62502: https://www.mathworks.com/matlabcentral/fileexchange/62502. Raw sequence reads are available on the NCBI SRA under accession number PRJNA380107: https://www.ncbi.nlm.nih.gov//bioproject/PRJNA380107.

## Author’s contributions

All four co-authors were involved in conception of the work; data analysis and interpretation; and drafting, critical review and final approval of the article. Data collection (TJK and JRP). Mathematical framework and numerical scheme (RDS and GDCS). TJK supervised by JRP. RDS was jointly supervised by JRP and GDCS.

## Acknowledgements

The work was supported in part by National Science Foundation Grant DMS 1121606 to GDCS, the Biomathematics Initiative at The College of William & Mary, and a W&M Summer Research grant to JRP.

## Author details

Department of Applied Science, The College of William & Mary, 23187, Williamsburg, VA, USA. ^2^Department of Biology, The College of William & Mary, 23187, Williamsburg, VA, USA.

